# Surface delivery quantification reveals distinct trafficking efficiencies among clustered protocadherin isoforms

**DOI:** 10.1101/2024.09.23.614616

**Authors:** Elizabeth J. May, Rachelle Gaudet

## Abstract

Proteins that transmit molecules and signals across the plasma membrane are crucial in cell biology because they enable cells to sense and respond to their surroundings. A major challenge for studying cell-surface proteins is that often they do not fold or traffic properly to the plasma membrane when produced in heterologous cells. We developed a strategy for quantifying surface localization from fluorescence microscopy images of surface-stained cells. Using clustered protocadherins, a protein family important for cell-cell recognition during neuronal development, we found that surface delivery levels vary among clustered protocadherin isoforms and between wildtype and engineered variants. Quantifying these differences provides evidence that *cis* dimerization is not tightly coupled to surface delivery for clustered protocadherins. This work establishes a generalizable framework for screening proteins and variants of interest for proper cell surface localization.

**Significance:** Surface proteins allow cells to interact with their environments, and their activities are often regulated by their delivery to and removal from the plasma membrane. We developed a strategy to quantitatively compare the surface delivery of proteins based on established epitope tag-based surface staining methods. Using natural and engineered variants of clustered protocadherins, cell-surface proteins essential for neuron development, we show that quantitative comparisons of surface trafficking levels facilitate the interpretation of mutational effects and can shed light on key regulatory mechanisms. We find that surface trafficking levels differ between variants and that, contrary to what was previously thought, a domain that inhibits surface delivery in some clustered protocadherins does so without directly relying on its protein-protein interface.

## Introduction

Clustered protocadherins (cPCDHs) are a large family of single-pass transmembrane proteins that function as adhesion and signaling molecules in brain development. They are primarily expressed in the nervous system (1–4) and their genetic deletion in mice is lethal at birth due to extensive neuronal cell death (5–8). At the cellular scale, cPCDH perturbations lead to a range of defects in neuronal cell morphology and connectivity, and their roles in dendrite complexity and self-avoidance are the most studied and best understood (9–14). These essential neuronal activities rely on homophilic cPCDH interactions at cell-cell contacts, so proper localization to the plasma membrane is a prerequisite for cPCDH function.

In mammalian genomes, including mice and humans, more than fifty genes encode a diverse set of paralogous cPCDH isoforms (4, 15, 16). Each isoform contains six extracellular cadherin (EC) domains, a transmembrane helix, and an intracellular region (**Figure 1A**). The cPCDH genomic locus includes clustered arrays of exons that encode the N-terminal extracellular domains and the first ∼100 amino acids of the intracellular region for each isoform. From there, the sequences differ only by subfamily: α-PCDHs and γ-PCDHs have α- and γ-specific C-terminal sequences encoded by shared exons, while β-PCDHs do not have shared exons and thus have shorter intracellular regions (**Figure S1A**) (3, 4, 16–18). At the cell surface, cPCDHs form strictly homophilic *trans* dimers between molecules on juxtaposed membranes and preferentially heterophilic *cis* dimers between isoforms on the same membrane (19–31). Defining these cPCDH interactions has led to a proposed mechanism wherein cPCDH oligomerization via both *cis* and *trans* dimers serves as a self-contact signal that ultimately leads to self-avoidance through an as-yet-unknown signaling pathway (**Figure S1B**) (20, 22, 23, 26–30, 32). Cell surface localization and *cis* and *trans* dimerization are all required for this mechanism.

**Figure 1.**
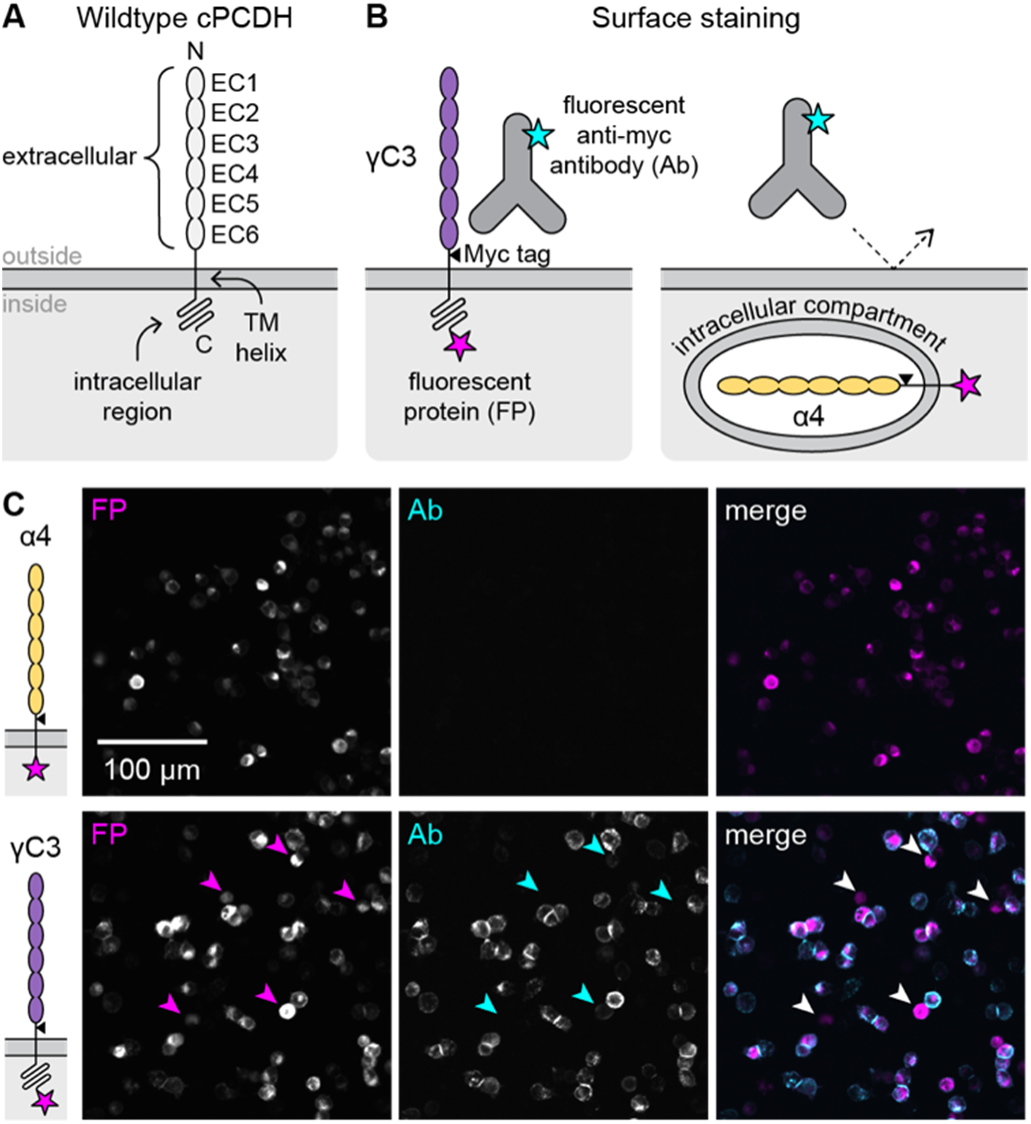
Surface staining by extracellular epitope detection of cPCDHs in 293T-ΔNC cells is the basis of surface delivery quantification. (A) Schematic showing the domain arrangement and topology of a generic cPCDH isoform. From N to C terminus: six extracellular cadherin (EC) domains are tethered to the membrane by a transmembrane helix (TM) followed by an unstructured intracellular region. (B) Diagram of experimental approach. cPCDH isoforms modified to contain an extracellular Myc epitope tag (triangle) and a C-terminal fluorescent protein (magenta star) are expressed in 293T-ΔNC cells. A fluorescent (cyan star) anti-Myc antibody recognizes the Myc tag to detect proteins that reach the cell surface, like γC3. The antibody does not bind proteins that are retained intracellularly, like α4, because it does not cross the plasma membrane. (C) Microscopy images of surface-stained control samples. Top row: An α-PCDH isoform, α4, is produced in transfected 293-ΔNC cells (signal in FP channel) but does not reach the cell surface (no signal in Ab channel). Bottom row: A C-type isoform, γC3, is produced in transfected 293T-ΔNC cells (signal in FP channel), and the anti-Myc antibody stains the periphery of cells with plasma membrane-localized protein (Ab channel). Arrowheads in γC3 images indicate examples where transfected cells have little to no Ab staining, highlighting the heterogeneity of surface trafficking.

In the brain and in cultured neurons, cPCDHs are not uniformly distributed on the cell surface; instead, they are largely intracellular, have a punctate distribution, and are often enriched at cell-cell contact sites (31, 33–39). When transfected into heterologous cells, α-PCDHs do not reach the cell surface, while β- and γ-PCDHs do (21). This differential trafficking has been attributed to *cis* dimerization (21, 26, 28, 40). The cPCDH *cis* interaction involves EC6 from one subunit and both EC5 and EC6 (EC5-6) from the other subunit, forming an asymmetric dimer (28). Previous work identified several conditions that induced α-PCDHs to go to the cell surface: (i) deletion of the EC6 domain, (ii) replacement of the α-PCDH EC6 domain with a β- or γ-PCDH EC6 domain, (iii) co-transfection with a full-length β- or γ-PCDH, or (iv) co-transfection with an EC5-6 fragment (but not an EC6-only fragment) of a β- or γ-PCDH (**Figure S1C**) (21). Disruptive mutations at *cis* interface positions in β- and γ-PCDHs also interfered with their surface delivery (26, 28). These observations led to the conclusion that *cis* dimerization is required for cPCDH surface delivery (21, 26, 28, 40), although the mechanism of this regulation remains unclear. In addition to the role of the extracellular domains described above, ubiquitination or phosphorylation of the γ-PCDH variable intracellular region may also play a regulatory role in surface trafficking (31, 38, 39, 41–44).

Several previous studies have included observations of cPCDH subcellular localization in heterologous cells (1, 19, 21, 33–35, 41, 45–50), but few have made quantitative surface delivery measurements (31, 49). To identify cPCDH features that influence their surface trafficking, we quantified cell surface localization using antibody detection of an extracellular epitope tag and developed generalizable metrics that summarize multiple aspects of diverse surface delivery phenotypes. We tested mutation-containing and domain-swapped α-PCDH variants to identify molecular features that enable α-PCDH cell surface trafficking. We found that while replacing the EC6 domain of an α-PCDH with EC6 of a β- or γ-PCDH increased surface localization compared to the wildtype α-PCDH, introducing γ-PCDH *cis* interface residues into an α-PCDH did not. EC6-swapped α-PCDH constructs also had lower surface delivery levels than wildtype β- and γ-PCDHs, and *cis* dimerization affinity did not correlate with wildtype β- and γ-PCDH surface trafficking levels. Our results indicate that features of EC6 beyond the *cis* dimer interface contribute to the surface delivery of α-PCDHs, and they suggest that *cis* dimerization-independent mechanisms also regulate cPCDH surface trafficking.

## Results

### Defining metrics for surface delivery quantification

To investigate the surface trafficking of cPCDHs, we pursued a quantitative approach that would facilitate comparisons of wildtype cPCDH isoforms and engineered variants. Surface staining by antibody detection of extracellular epitope tags is a common approach to assess a protein’s surface localization that has previously been applied to cPCDHs in heterologous cell lines and cultured neurons (21, 34, 48–51). We used a derivative of the 293T cell line that lacks the adhesion protein N-cadherin (293T-ΔNC); these cells are adherent but detach easily and can be used for cell aggregation assays (52). Using cPCDH constructs containing an extracellular Myc tag and an intracellular fluorescent protein (FP; **Figure S1A**), we added fluorescent anti-Myc antibodies (Ab) to transfected cells and collected fluorescence microscopy images. We first compared two wildtype cPCDH isoforms known to have different surface trafficking behaviors: α4, which does not traffic to the plasma membrane, and γC3, which does (**Figure 1B**) (21). The FP served as a transfection marker, and transfection efficiencies were similar across samples (**Figure S1D**). Notably, the cellular distribution of the FP signal was indistinguishable between α4 and γC3, although the overall FP intensity was different. Ab signal was undetectable in the α4-transfected sample (**Figure 1C**), confirming the non-surface localization of α4. As expected, the Ab stained the cell periphery and cell-cell junctions at contacts between transfected cells in the Myc-tagged γC3 sample (**Figure 1C**).

Although the overall surface delivery behaviors of α4 and γC3 were apparent in our images, we noticed that some γC3-expressing cells had very little or no measurable surface staining (**Figure 1C, arrowheads**). We therefore devised metrics to account for the phenotypes of individual cells and evaluate overall surface delivery based on the distribution of cellular phenotypes. For each sample, we collected both fluorescence images in the FP and Ab channels and brightfield images (**Figure 2A**). We used automated segmentation in brightfield to identify cells and calculated the mean FP and Ab signal per cell (*FP_cell_* and *Ab_cell_*) from the fluorescence channels (**Figure 2A**). We identified transfected cells using a threshold *FP_cell_* value determined by an untransfected control (**Figure 2B**). We then generated surface stain histograms of the *Ab_cell_* values of transfected cells to visualize the spread of surface delivery phenotypes within samples (**Figure 2C**).

**Figure 2.**
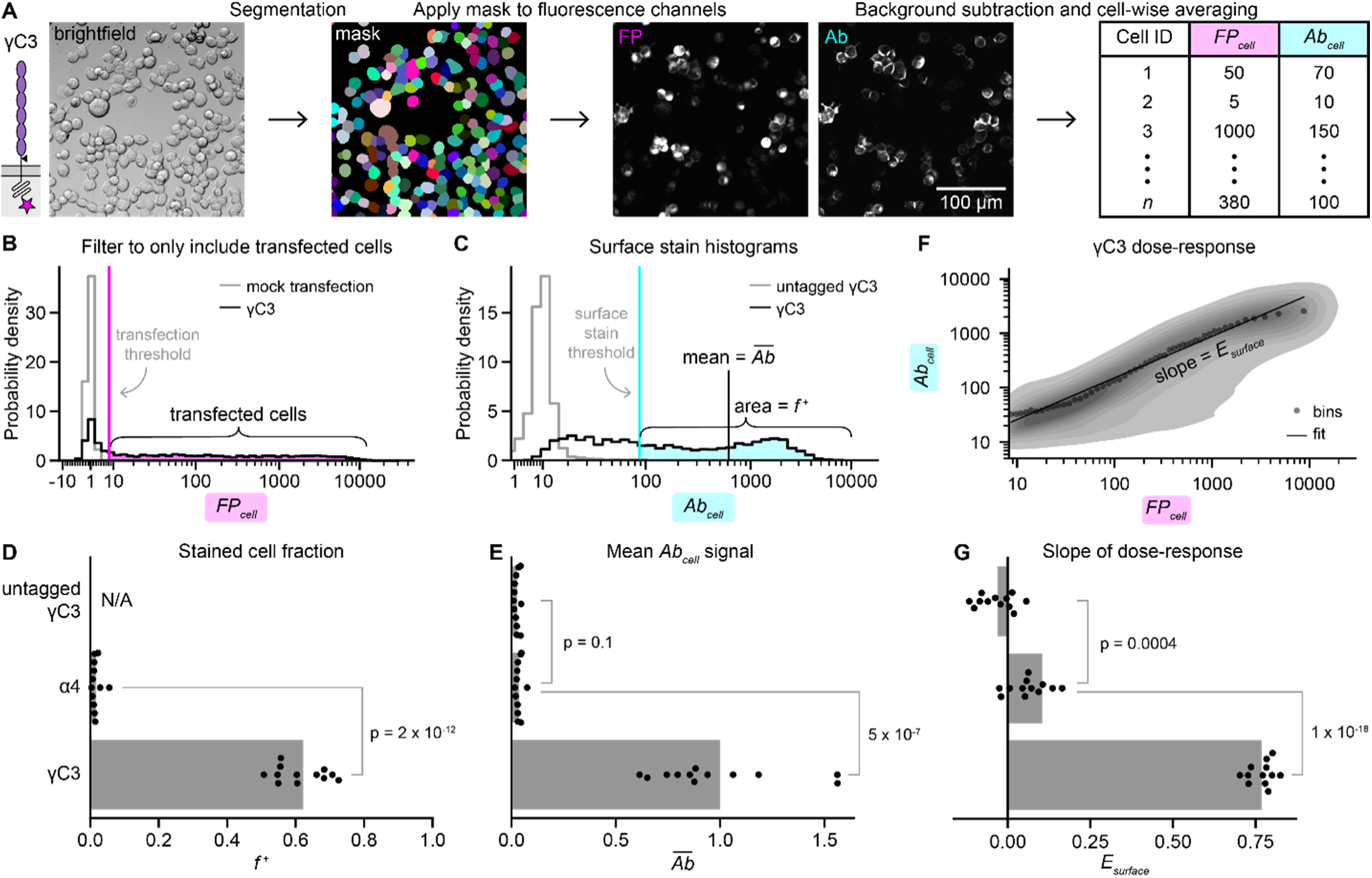
Quantification reveals the heterogeneity of surface trafficking across cells. (A) Initial image processing steps for an example sample, γC3. Automated cell segmentation using the brightfield channel from a particular field of view produces a mask that assigns each pixel in the field of view to a cell (or background). To perform background subtraction, we applied the mask to the FP and Ab images from the same field of view to assign background pixels, and we subtracted the average background pixel values across the image. Cell-wise averaging of background-subtracted pixel intensities produces *FP_cell_* and *Ab_cell_* values for each of *n* cells in the field of view. (B) A histogram of *FP_cell_* values, where *FP_cell_* is a measure of cellular protein levels of the transfected cPCDH. The data are filtered to only include transfected cells (magenta highlight) using a transfection threshold (magenta vertical line) determined using a sample of mock-transfected cells. (C) A histogram of the filtered *Ab_cell_* values, where *Ab_cell_* a measure of surface abundance of the transfected cPCDH. The mean of the entire *Ab_cell_* distribution, *Ab^-^* (black vertical line), corresponds to the average surface stain signal for all transfected cells in the sample. The fraction of Ab-stained cells, *f^+^* (cyan highlight), is calculated using a threshold *Ab_cell_* value (cyan vertical line) determined using cells transfected with an untagged version of γC3. (D-E) Graphs of *f^+^* (D) and *Ab^-^*(E) for untagged γC3, α4, and γC3. In panel E, *Ab^-^* values are normalized to the mean *Ab^-^* of Myc-tagged γC3. (F) Dose-response plot of the γC3 data. The contour plot displays the density of *FP_cell_* and *Ab_cell_* values for each cell. The data were binned by *FP_cell_* and the means of each bin are shown (gray points). The bins were scaled to maintain the same number of cells in each bin, which resulted in approximately log-scaled bins. We define *E_surface_* as the slope of a linear fit to the data (black line). (G) Graph of *E_surface_* for untagged γC3, α4, and γC3. In panels D, E, and G, points are values for biological replicates (*n* = 12), and bars show the mean values. P-values were calculated using a one-tailed Welch’s t-test.

Using a γC3 control construct lacking the extracellular Myc tag (**Figure 2C**, gray histogram), we determined a threshold *Ab_cell_* value (**Figure 2C**, cyan vertical line) and calculated the fraction of transfected cells with measurable surface staining, *f^+^*(**Figure 2C**, cyan shaded region). For γC3, *f^+^* was 0.62 ± 0.07 (mean ± SD), while for α4 *f^+^* was close to zero (0.01 ± 0.01; **Figure 2D**). We also calculated the mean *Ab_cell_* value, *Ab^-^*, which is proportional to the average cell-surface cPCDH concentration for a given construct (**Figure 2C**, black vertical line). As expected, *f^+^* and *Ab^-^* for γC3 were significantly higher than for α4 (p = 2×10^-12^ for *f^+^* and p = 5×10^-7^ for *Ab^-^*; **Figure 2E**).

We next investigated how the amount of protein on the cell surface depends on expression level. Taking advantage of the variation in expression generated by transient transfection and reported by our transfection marker, we constructed dose-response curves, which exhibited approximately power law scaling when plotted linearly (**Figure S1E**). To group cells by expression level, we binned by *FP_cell_* and calculated the mean *Ab_cell_* for each bin (**Figure 2F**; gray dots). On a log scale, the slope of a linear fit to these data represents a surface delivery “efficiency” (*E_surface_*) that is related to the fraction of protein produced that localized to the cell surface. By accounting for protein expression level, *E_surface_* facilitates meaningful comparisons of surface trafficking between constructs even when expression levels vary. *E_surface_* was positive for γC3, indicating that cells making more γC3 protein overall had more γC3 on the surface than cells with lower γC3 expression levels, whereas *E_surface_* for α4 was close to zero, showing that there was little to no surface staining of α4-transfected cells at any expression level (**Figure 2G**).

### Quantification can distinguish different surface delivery levels

To validate our quantification strategy and investigate the relationship between cPCDH *cis* dimerization and surface delivery, we focused on the *cis* interface domain EC6. We tested EC6-swapped α4 constructs (α4-γB6_EC6_, α4-β17_EC6_, and α4-γC3_EC6_), and their wildtype counterparts (γB6, β17, and γC3) for surface delivery in 293T-ΔNC cells (**Figure 3A**). We chose these variants because they were all previously shown to mediate cell aggregation in K562 cells (21, 22, 26). Cells transfected with EC6-swapped α4 variants had dim, punctate Ab signal (**Figure S2A**), often at contact points between transfected cells (**Figure S2B**). Wildtype β- and γ-PCDHs had more uniform surface staining with bright signal at cell-cell contacts, and β17 and γC3 were brighter than γB6 (**Figure S2A**). These samples thus presented a range of surface delivery phenotypes.

**Figure 3.**
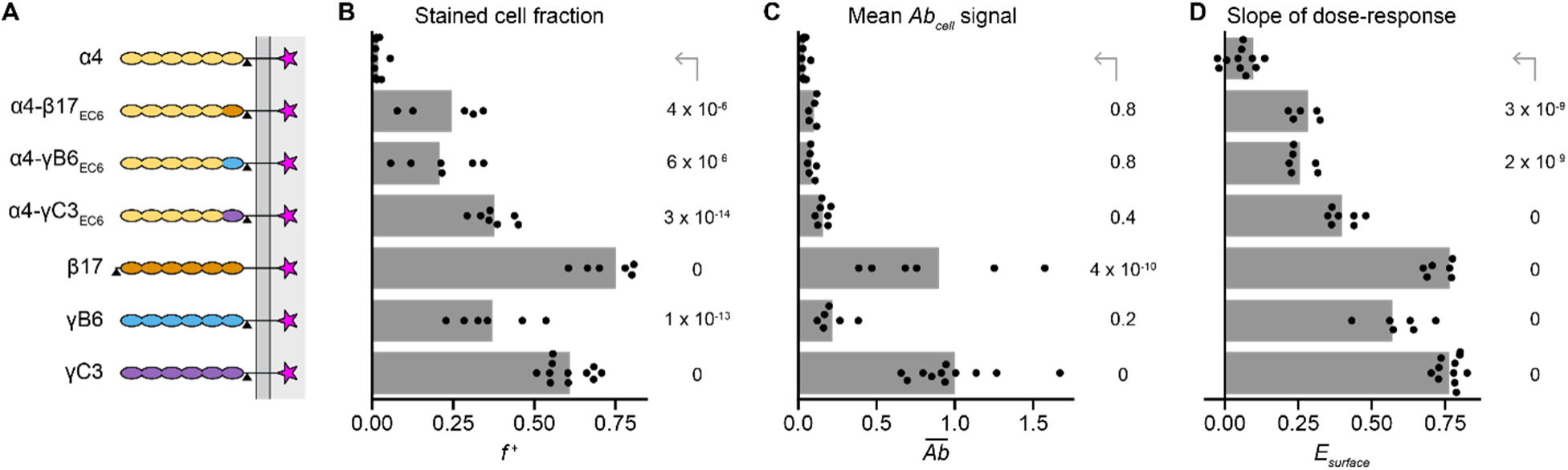
Quantification can distinguish samples with intermediate surface delivery phenotypes. (A) Schematics of tested constructs; domains are color-coded by originating isoform. (B-D) Quantification of *f^+^* (B), *Ab^-^* (C), and *E_surface_* (D) for the constructs shown in A. In B, *Ab^-^* values are normalized to the mean *Ab^-^* of the positive control construct, Myc-tagged γC3. Points are biological replicates (*n* = 11 for α4 and γC3; 5 for α4-β17_EC6_; 6 for α4-γB6_EC6_, β17, and γB6; and 7 for α4-γC3_EC6_), and bars are mean values across replicates. P-values for each sample compared to α4 (top row) were calculated using a one-tailed Dunnett’s test.

All the *Ab_cell_* histograms at least partially overlapped with the untagged γC3 distribution, showing that every sample contained a subpopulation of transfected cells without any surface stain signal (**Figure S3**). The EC6-swapped α4 variants had *f^+^*between 0.2 and 0.4, while *f^+^* for the wildtype β and γ isoforms was generally higher and ranged from 0.4 to 0.7 (**Figure 3B**). *Ab^-^* values for the wildtype β17 and γC3 isoforms were significantly higher than *Ab^-^* for wildtype α4 (p ≤ 4×10^-10^; **Figure 3C**). The *Ab^-^* metric emphasized the ultra-bright surface staining of some cells in the β17 and γC3 samples (**Figure S3**), indicating that β17 and γC3 traffic to the cell membrane particularly well for cPCDHs. Compared to α4, *E_surface_* was positive and significantly higher for all other constructs (p ≤ 3×10^-9^), and *E_surface_* was highest for the wildtype β and γ isoforms (**Figure 3D**).

Overall, our quantification indicates that surface levels of the EC6-swapped variants were higher than wildtype α4 in 293T-ΔNC cells, in agreement with previous findings that the same variants mediated cell aggregation in K562 cells (21, 22, 26). Furthermore, our quantification revealed that the wildtype isoforms (γB6, β17, and γC3) generally trafficked better than the engineered α4 variants (α4-γB6_EC6_, α4-β17_EC6_, and α4-γC3_EC6_).

Finally, to validate our use of the 293T-ΔNC cell line for surface trafficking measurements, we also measured surface delivery in K562 cells. K562 cells grow in suspension, so we measured *FP_cell_* and *Ab_cell_* using flow cytometry. Using α4, α4-γC3_EC6_, and γC3, we confirmed that the trends we observed were consistent across cell types and measurement modalities (**Figure S4**). As in 293T-ΔNC cells, both α4-γC3_EC6_ and γC3 were significantly higher than α4 in K562 cells (p = 5×10^-11^ and 0 for *E_surface_*, respectively; **Figure S4B**), and γC3 was significantly higher than α4-γC3_EC6_ (p = 5×10^-7^ for *E_surface_*; **Supplementary Dataset S1**).

### The γ-PCDH *cis* interface is insufficient to enable α-PCDH surface delivery

We next explored the link between cPCDH *cis* dimerization and surface delivery to investigate why α-PCDHs do not localize to the cell surface. Previous studies used a loss-of-function approach—introducing disruptive mutations into the *cis* interface of representative β- and γ-PCDHs—to show that some, but not all, *cis* interface mutations prevented cell aggregation in K562 cells (26, 28). This finding supported the conclusion that *cis* dimerization is necessary for cPCDH surface delivery (21, 26, 28, 40). To clarify the role of *cis* dimerization in α-PCDH surface delivery, we took a complementary gain-of-function approach—introducing β- and γ-PCDH amino acids into an α-PCDH at differentially conserved *cis* interface positions—to test whether *cis* interface residues are sufficient to increase α-PCDH surface delivery.

To identify differentially conserved *cis* interface positions, we aligned all wildtype mouse cPCDH isoforms using their extracellular sequences (EC1-6) and divided the sequences into two alignments according to their cell aggregation behavior as reported by Thu et al. (21): cell surface trafficking isoforms (β- and γ-PCDHs, αC2, γC3, and γC5) and non-trafficking isoforms (α-PCDHs, αC1, and γC4). We calculated the amino acid frequencies at each position in EC6 separately for the two alignments and compared the resulting distributions using the Kullback-Leibler divergence (*KL_div_*; **Figure 4A**). *KL_div_* scores are high for positions that are conserved within each alignment but differ between the two alignments, and low for positions that are either variable or conserved across both alignments.

**Figure 4.**
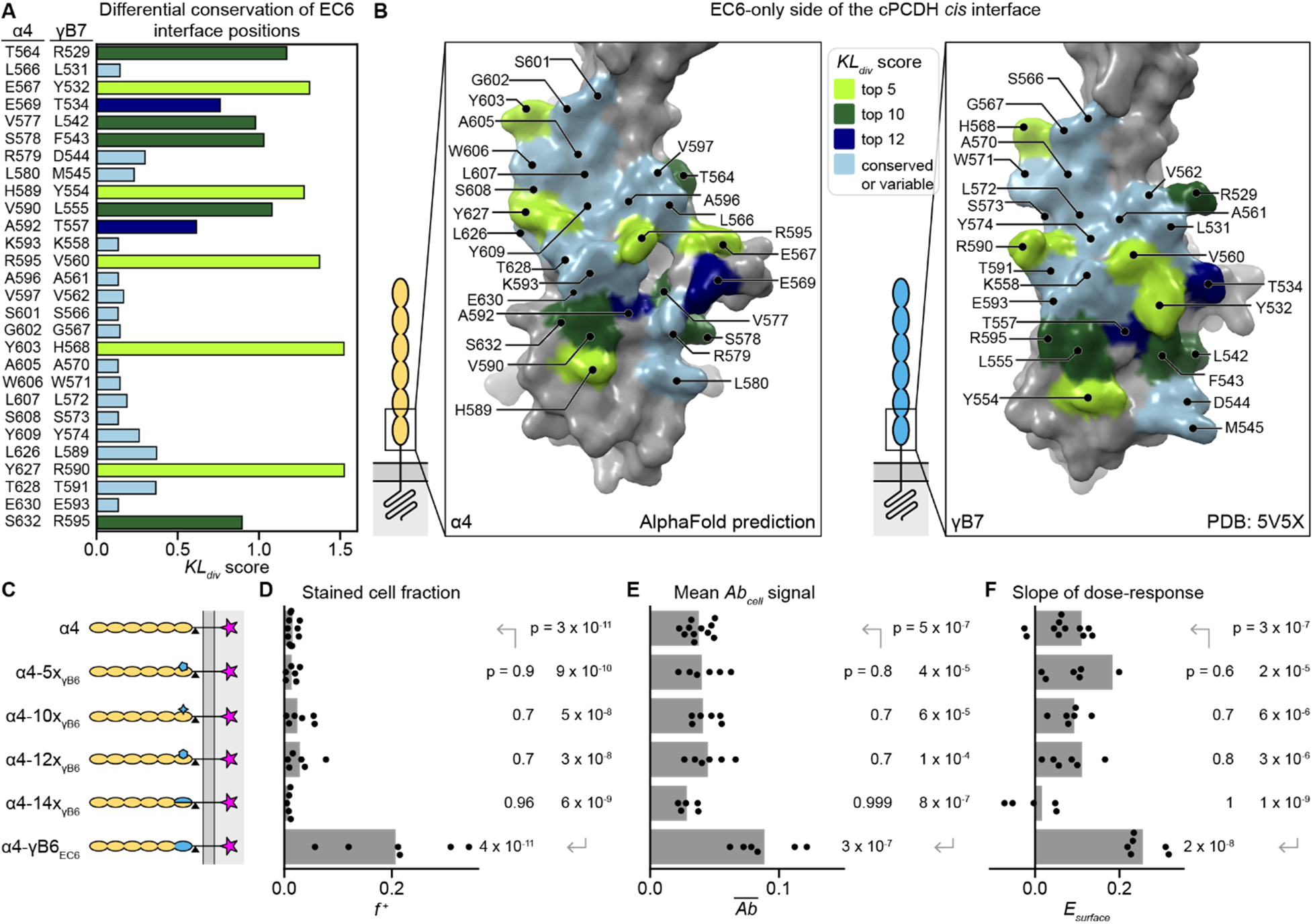
The EC6 *cis* interface of γB6 is insufficient for surface delivery of α4. (A) Differential conservation for all EC6 *cis* interface positions quantified using Kullback-Leibler divergence (*KL_div_*) score, which compares the conservation of a position among isoforms that traffic to the cell surface to that position’s conservation among isoforms that do not reach the surface. The color of the bar indicates how positions were grouped by KL_div_ score for chimeric constructs (see legend in panel B). (B) Surface representations of the EC6 *cis* interface for α4 (left; structure prediction from AlphaFold Protein Structure Database [accession code: AF-O88689-F1-v4] (53)) and γB7 (right; experimentally-determined structure [PDB: 5V5X] (28)). *Cis* interface residues are colored according to their *KL_div_* score categories (legend top middle). (C) Schematics of the chimeric α4 constructs quantified in panels D-F. (D-F) Quantification of *f^+^* (D), *Ab^-^* (E), and *E_surface_* (F) for the constructs shown in panel C. In E, *Ab^-^* values are normalized to the mean *Ab^-^* of the positive control construct γC3 (not shown in the figure). Points are biological replicates (*n* = 11 for α4; 5 for α4-14x_γB6_; and 6 for α4-5x_γB6_, α4-10x_γB6_, α4-12x_γB6_, and α4-γB6_EC6_), and bars are replicate means. P-values for each sample compared to α4 were calculated using a one-tailed Dunnett’s test. P-values for each sample compared to α4-γB6_EC6_ were calculated using a two-tailed Dunnett’s test.

We mapped the *KL_div_* scores of EC6-side *cis* interface residues onto the γB7 *cis* dimer structure (28) and the predicted α4 model from the AlphaFold Protein Structure Database (53) (**Figure 4B**).

In general, the residues in the top and center of the *cis* interface are strictly conserved among all cPCDHs, while differentially conserved positions are at the sides and bottom of the interacting surface, as previously noted (28). Several of the most differentially conserved positions in the *cis* interface have different sidechain properties (e.g., R595 in α4 versus V560 in γB7, or Y627 in α4 versus R590 in γB7; **Figure 4B**). To test whether these differences underlie subfamily-specific surface delivery, we introduced conserved β- and γ-PCDH *cis* interface residues into the corresponding positions of an α-PCDH.

We continued using α4 as a representative non-trafficking α-PCDH isoform. To make α4 more likely to *cis* homodimerize, we introduced residues from a γB-subfamily isoform because surface plasmon resonance measurements of cPCDH *cis* heterodimers suggested that γB-subfamily cPCDHs are the most amenable to participating as either the EC6-only or EC5-6 sides of the *cis* interface (29). We chose γB6 because it is highly similar to γB7, the only isoform for which a *cis* dimer structure is available (28), with 97% sequence identity in EC6. Of the 28 EC6 interface residues, fourteen differ between α4 and γB6. We mutated the top five (α4-5x_γB6_), top ten (α4-10x_γB6_), and top twelve (α4-12x_γB6_) most differentially conserved amino acids in α4 to the corresponding γB6 amino acid, as well as making all possible EC6 interface mutations (α4-14x_γB6_; **Table S1; Figure 4C**). Contrary to our hypothesis that these would be gain-of-function mutations, none of the mutation-containing chimeras showed improved surface delivery compared to wildtype α4, even though the EC6 domain-swapped construct α4-γB6_EC6_ did (**Figure 4D-F**). To verify that the mutations did not destabilize the protein, we co-expressed α4 and the chimeric constructs with untagged γA11 (**Figure S5A**). Wildtype α4 and all the mutation-containing variants except α4-14x_γΒ6_ were detected on the surface to similar extents in the co-transfection experiment (p ≤ 0.03 comparing *E_surface_* of each co-transfection experiment to α4 expressed on its own; **Figure S5B-D**). Although α4-14x_γΒ6_ was expressed at similar levels to surface-trafficking constructs such as α4-γB6_EC6_ (**Figure S5E**), it is possible that α4-14x_γΒ6_ does not fold properly, and that at least some α4 EC6 *cis* interface positions are important for the protein’s stability.

Overall, we found that changing differentially conserved EC6 *cis* interface positions of α4 to match γB6 was insufficient to increase surface delivery compared to wildtype α4. This was surprising because α-PCDHs were previously thought to not reach the cell surface because they were unable to form *cis* homodimers (28), which implied that the *cis* dimer interface in the γB6 EC6 domain was responsible for the surface delivery of α4-γB6_EC6_. In contrast, our results suggest that EC6 features away from the *cis* dimer interface explain the domain-swap-rescue phenotype for α-PCDHs.

### Surface levels of EC6-swapped α-PCDHs are lower than wildtype β- and γ-PCDHs

To further explore the EC6 features that explain why a domain-swap can rescue surface delivery of α-PCDHs, we generated an expanded set of EC6-swapped α4 constructs using isoforms from the different cPCDH subfamilies (**Table S1; Figure 5A**). We also tested the corresponding wildtype β or γ isoforms, and an α4 variant lacking EC6 altogether (α4ΔEC6; **Figure S6A**). The α4ΔEC6 construct showed a significantly higher *E_surface_* than wildtype α4 (p = 2×10^-5^; **Figure 5B**), and most EC6-swapped α4 constructs trafficked like α4ΔEC6 (p ≥ 0.2 for *E_surface_* for all EC6-swapped α4 constructs compared to α4ΔEC6), except for α4-γC3_EC6_, which had higher *E_surface_* than α4ΔEC6 (p = 0.0002; **Figures 5B and S6B-C**). This suggests that surface delivery is enabled by the absence of the α4 EC6 domain, not the presence of a β- or γ-PCDH EC6 domain, except for the γC3 EC6 domain possibly further enhancing surface trafficking.

**Figure 5.**
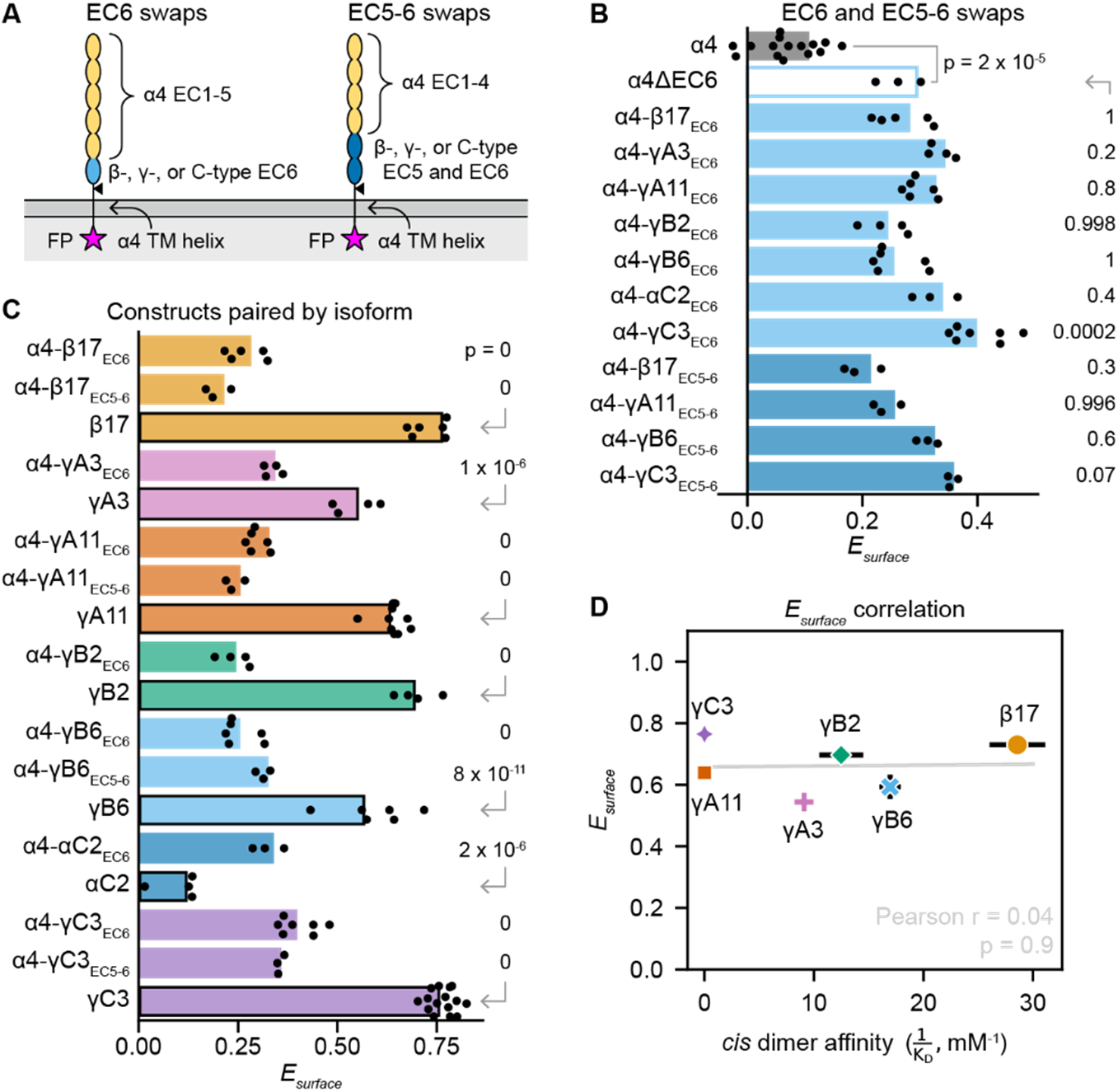
Removing or replacing EC6 of α4 increases its surface delivery, but not to wildtype β- and γ-PCDH levels. (A) Schematics of the types of constructs compared in B-D. EC6-swapped variants have EC1-5 and the transmembrane helix of α4, and EC6 of a different isoform. EC5-6 swaps have EC1-4 and the transmembrane helix of α4, and EC6 of a different isoform. (B) Graph comparing *E_surface_* for wildtype α4 (gray bar) to α4 variants with EC6 deleted (unfilled light blue bar), EC6 swapped (lighter blue bars), or EC5-6 swapped (darker blue bars). Points are biological replicates (*n* = 15 for α4 and γC3; 3 for α4ΔEC6, α4-αC2_EC6_, α4-β17_EC5-6_, α4-γA11_EC5-6_, α4-γΒ6_EC5-6_, and γC3_EC5-6_; 4 for α4-γA3_EC6_ and α4-γB2_EC6_; 5 for α4-β17_EC6_; 6 for α4-γA11_EC6_ and α4-γB6_EC6_; and 7 for α4-γC3_EC6_), and bars are replicate means. The p-value for α4ΔEC6 compared to α4 was calculated using a one-tailed Welch’s t-test; p-values for domain-swapped samples compared to α4ΔEC6 were calculated using a two-tailed Dunnett’s test. (C) Graph of *E_surface_* comparing constructs grouped by isoform, with bars color-coded by isoform. Bars without borders are α4 variants (EC6 or EC5-6 swaps), while bars with black borders are wildtype cPCDHs. Points are biological replicates (*n* = 15 for α4 and γC3; 3 for α4-αC2_EC6_, α4-β17_EC5-6_, α4-γA11_EC5-6_, γB6, α4-γΒ6_EC5-6_, and γC3_EC5-6_; 4 for αC2, γA3, α4-γA3_EC6_, γB2, and α4-γB2_EC6_; 5 for α4-β17_EC6_; 6 for β17, α4-γA11_EC6_, γB6, and α4-γB6_EC6_; 7 for α4-γC3_EC6_; and 10 for γA11), and bars are replicate means. P-values for each α4 variant compared to the corresponding wildtype isoform were calculated using Tukey’s test. (D) Scatter plot of wildtype cPCDH *cis* dimerization affinity versus surface delivery, as measured by *E_surface_*. See Figure S6F for affinity values used in this plot. Points are shape- and color-coded by isoform. The Pearson r statistic and its associated p-value reflect the degree of correlation.

Our results agree with previous observations in K562 cells suggesting that the α-PCDH EC6 domain inhibits surface trafficking, including that α4ΔEC6 trafficked to the cell surface and mediated cell aggregation, and that replacing γC3’s EC6 with α4’s EC6 interfered with surface trafficking and cell aggregation (21). Because co-expression of α-PCDHs with β- or γ-PCDHs led to α-PCDH surface trafficking (21, 22, 26, 28), a finding which we reproduced here in 293T-ΔNC cells (**Figure S5**), it was previously inferred that the α-PCDH *cis* dimer interface contained a negative regulator that was responsible for the inhibitory activity of EC6, and that *cis* heterodimerization with a β- or γ-PCDH could mask this inhibitory retention signal. Together with our observation that extensive mutagenesis of the α4 EC6 *cis* dimerization interface did not increase surface trafficking, our quantitative surface-trafficking results instead suggest that the α-PCDH EC6 inhibits surface trafficking through an unknown mechanism that does not require the *cis* dimer interface.

We next compared the surface delivery of the EC6-swapped α4 variants to that of the originating wildtype isoforms. For all constructs except αC2, the EC6-swapped variant trafficked less than the corresponding wildtype (p ≤ 1×10^-6^ for *E_surface_*) (**Figure 5C, Figure S6D-E**). αC2 is part of the α-PCDH genomic cluster but is phylogenetically distinct from the α-PCDHs, like the other C-type isoforms (16). Wildtype αC2 trafficked poorly, with *f^+^*, *Ab^-^*, and *E_surface_* similar to α4 (p = 0.2 for *E_surface_*; **Figure S4**), while α4-αC2_EC6_ trafficked better than α4 (p = 1×10^-14^ for *E_surface_*; **Figure 5B**), similar to the other EC6-swapped variants (**Figure 5B-C, Figure S6B-E**). This suggests that the αC2 EC6 does not share α-PCDH EC6’s inhibitory feature. We note that in previous work, the same αC2 construct was detected on the cell surface and mediated cell aggregation in K562 cells (21, 22), albeit at low levels. In our hands, full-length and cytoplasmic region-deleted αC2 trafficked as poorly as α4 in both K562 and 293T-ΔNC cells (**Figure S4**). However, the overall trend that α4ΔEC6 and EC6-swapped α4 variants did not traffic as well as wildtype β- and γ-PCDHs suggests that additional features beyond EC6 also contribute to determining α-PCDH surface delivery levels.

As cPCDH *cis* dimerization is asymmetric, with EC6 of one subunit binding EC5 and EC6 of the other subunit, we wondered whether α4 constructs with both EC5 and EC6 swapped would traffic better than EC6-swapped α4 variants. We generated EC5-6-swapped α4 variants that represented each cPCDH subfamily: α4-γA11_EC5-6_, α4-γB6_EC5-6_, α4-β17_EC5-6_, and α4-γC3_EC5-6_ (**Table S1; Figure 5A, Figure S6A**). The EC5-6-swapped α4 constructs showed no improvement over their EC6-swapped counterparts; rather, they were also similar to α4ΔEC6 (p ≥ 0.07 for *E_surface_*; **Figure 5B**). Therefore, features of EC6, but not EC5, contribute to determining α-PCDH surface delivery levels.

### Surface delivery levels are independent of *cis* dimerization affinity

Previous studies using *cis*-dimer-disrupting mutations in β- and γ-PCDHs revealed that the *cis* dimer interface is necessary for β- and γ-PCDH surface delivery (26, 28). We investigated the relationship between *cis* binding affinity and surface delivery levels in wildtype β- and γ-PCDH isoforms. We looked at whether our quantitative surface trafficking measurements of wildtype β- and γ-PCDHs correlate with published *cis* homodimerization affinity values (or that of the closest paralog for which an affinity has been reported (22, 26–29); **Figure S6F)**. If *cis* dimerization is the primary determinant of surface trafficking levels, we expect isoforms that dimerize with higher affinity to traffic better than lower affinity isoforms. We found no convincing correlation between affinity and any of our surface delivery metrics (**Figure 5D, Figure S6G-H)**. Overall, *cis* dimerization affinity was a poor predictor of surface delivery levels for β- and γ-PCDH isoforms, suggesting that it is not the sole determinant of β- and γ-PCDH export.

## Discussion

In heterologous expression systems, surface delivery is a useful metric to validate proper surface localization for proteins of interest, to account for differences in surface protein levels, and to identify protein modifications that disrupt trafficking. We developed a robust method to quantify the surface delivery of plasma membrane proteins using epitope-tag-based surface staining. Our metrics describe three axes of variation common to any surface staining experiment: the fraction of cells with detectable levels of surface protein, relative surface protein abundance, and relative surface trafficking efficiency. We show that our quantification strategy is compatible with data acquired either by fluorescence microscopy or flow cytometry, and our scripts are available for researchers to modify or apply directly to their own datasets. To validate our approach, we applied it to cPCDHs and found that the quantification recapitulated known subfamily-specific trafficking trends observed qualitatively via K562 cell aggregation assays: β- and γ-PCDHs reach the plasma membrane, while α-PCDHs do not (21, 22, 26, 34, 45). We also found that while removing EC6 from PCDHα4 enabled its surface localization, replacing α4’s EC6 or EC5-6 with β- or γ-PCDH domains imparted no additional increase in surface trafficking. Our measurements further showed varying trafficking levels among wildtype and engineered cPCDH isoforms, enabling analyses that tested the relationship between cPCDH *cis* dimerization and surface delivery.

*Cis* dimerization is essential for cPCDH function in self-versus non-self recognition (54), and previous studies investigating why α-PCDHs do not traffic to the cell membrane in heterologous cells concluded that *cis* dimerization is required for cPCDH surface localization (21, 26, 28). Two key observations supported this conclusion: (1) co-expressing an α-PCDH with a β- or γ-PCDH enabled surface localization of the α-PCDH (21, 26, 28, 54), and (2) dimer-disrupting mutations in γ-PCDHs, including mutations introducing α-specific residues, prevented their surface trafficking (26, 28). We were therefore surprised to find that incorporating γ-PCDH *cis* interface residues into an α-PCDH was insufficient to enable surface delivery. We also found that the *cis* dimerization properties of our constructs—EC6 identity for domain-swapped α-PCDH variants and binding affinity for wildtype isoforms—were not correlated with surface delivery levels. As a whole, while previous work indicated that *cis* dimerization is necessary for cPCDH surface delivery (21, 26, 28, 54), our data suggest that at least for α-PCDHs, *cis* dimerization is not sufficient for surface trafficking, and that *cis* dimerization is not the sole determinant of cPCDH export levels. Interestingly, this shifted interpretation re-opens the question of whether α-PCDHs can form *cis* homodimers in cells, as the conclusion that they cannot has been based on their surface trafficking behavior in non-neuronal cells.

Previous research with α4ΔEC6 in K562 cells led to the conclusion that the α-PCDH EC6 domain inhibits surface delivery (21). In replicating these findings in 2963T-ΔNC cells, we showed that inhibition by the α-PCDH EC6 domain was consistent across cell types. Our finding that replacing EC6 in α4 with EC6 from a β-, γ-, or C-type cPCDH did not improve surface delivery compared to α4ΔEC6 also substantiates that idea, and it emphasizes that the lack of the α-PCDH EC6 matters more for the surface delivery of these variants than particular features of β- and γ-PCDH EC6 domains. Furthermore, our α4-γB6 *cis* interface chimeras showed that the *cis* interface is not the α-PCDH EC6 region that inhibits surface trafficking, and our affinity-versus-surface-delivery analysis suggests that *cis* dimerization is not the sole determinant for β- and γ-PCDH export.

Investigations into the cellular mechanisms of cPCDH trafficking suggest potential alternative models for how surface delivery might be controlled independently of *cis* dimerization. The cPCDH EC domains are O-mannosylated, an unusual type of glycosylation in metazoans catalyzed by four glycosyltransferases dedicated to cadherin superfamily proteins, including cPCDHs (55).

As protein glycosylation and folding are tightly linked quality control mechanisms in the secretory pathway, differences in glycosylation site motifs or glycosyltransferase interactions could determine the trafficking efficiency of a particular cPCDH isoform. Although we focused on extracellular interaction interfaces here, a motif in the variable intracellular region of γ-PCDHs has been shown to also regulate their trafficking in a ubiquitinylation- and phosphorylation-dependent manner in 293T cells (38, 39, 43, 44). Finally, correlative light and electron microscopy studies have identified specialized intracellular compartments that appear when γ-PCDHs, but not α-PCDHs, are overexpressed in 293T cells, raising the possibility that α- and γ-PCDHs may be trafficked by distinct intracellular pathways (39, 41). Quantification of surface delivery with the tools presented here will facilitate future investigations of the complex cellular regulation of cPCDH surface delivery in both heterologous cells and neurons.

In the cPCDH field, cell aggregation assays have played a crucial role in deciphering the molecular rules of cPCDH-mediated cell recognition (20–22, 26, 28, 54). Perhaps most importantly, sorting occurs in cell aggregation assays, with cells preferring to bind others with completely matching sets of cPCDH isoforms, the same scenario that leads to self-recognition and avoidance in neurons (21, 22, 54). Notably, Wiseglass and colleagues recently demonstrated that *cis* dimerization is essential for cPCDH-mediated sorting in cell aggregation assays (54). In these experiments, the extent of cell aggregation and sorting depended on relative isoform expression levels (20, 21, 54). Direct measurements of surface trafficking could provide insight into the relative surface concentrations of different isoforms in co-transfection experiments and distinguish how each isoform contributes to the energetics of competitive binding.

A limitation of studying membrane trafficking regulation in heterologous cells is that regulation may differ in the native context. Overexpression may overwhelm aspects of the secretory pathway, which could obscure which regulatory features are crucial for export. Of our three surface trafficking metrics, the mean *Ab_cell_* signal is highly dependent on expression level, while the threshold-based stained cell fraction metric, *f^+^*, is less so. Both of these metrics should be evaluated cautiously in contexts where expression levels vary, such as in transient transfections or when drawing comparisons between different proteins. In contrast, the *E_surface_* metric directly accounts for expression levels by measuring how strongly the amount of surface protein depends on the amount of protein expressed, making it independent of expression level.

Beyond cPCDHs, our method provides a general-purpose, quantitative way to screen plasma membrane proteins and variants of interest for surface delivery or secretion via either microscopy or flow cytometry. For attached cells, tissues, and adhesion protein studies, microscopy is preferable to avoid disrupting surface proteins when detaching cells from the culture vessel and each other. For suspended cells, flow cytometry has the advantage of being extremely high-throughput and therefore can detect rare events. Our surface delivery summary metrics are compatible with either approach. Cell surface proteins including ion channels, G-protein coupled receptors, and transporters represent highly important drug targets, and their binding pockets, gating and regulatory mechanisms are often mapped through structure determination and mutational scans (56–63). Membrane protein instability and mis-localization are common in heterologous expression systems (61, 64), and few screening studies validate loss-of-function hits by evaluating surface delivery (57, 59, 60, 63). Implementing quantitative surface delivery analysis in combination with activity screens will facilitate clearer interpretation of these experiments and improve investigations of surface protein functions.

## Materials and Methods

### Differential conservation sequence analysis

The sequences of all mouse cPCDHs were manually gathered from UniProt (release 2023_05) (65), aligned using MUSCLE v.5.1 (66), and truncated to EC6 (Geneious Prime 2023.1.2 [http://www.geneious.com]). The subsequent alignment manipulations were performed in Python 3 using bioviper v.0.20 (67). The alignment was separated into two sets of sequences: α-PCDHs, αC1, and γC4 in one set, and β-PCDHs, γ-PCDHs, αC2, γC3, and γC5 in the other set. For each sub-alignment, the amino acid frequencies were calculated and the frequency distributions at each position were compared between the two alignments using the entropy function in scipy v.1.11.4 (68) to calculate the Kullback-Leibler divergence, which we refer to as the *KL_div_* score.

### Molecular cloning

The wildtype-like constructs used in this study are illustrated in **Figure S1A**. The myc-tagged α4, α4ΔEC6, β17, αC2, and γC3 plasmids, and the untagged γC3 plasmid, were provided by Tom Maniatis’s lab (21). The α4 and α4ΔEC6 proteins, and all α4 variants used in this study, lacked the C-terminal intracellular region. The β17 construct lacked its C-terminal 25 intracellular residues, and αC2, γC3, and untagged γC3 were full length. The γA3, γA11, γB2, and γB6 constructs lacked the C-terminal intracellular region. All derived plasmids were cloned into the pmax-mCherry backbone of the α4 plasmid using Gibson assembly (Gibson Assembly Master Mix, NEB E2611S). For derived constructs including the myc tag, the tag was positioned in the stalk region between EC6 and the transmembrane helix (**Figure S1A**), equivalently to previously validated constructs (21). Inserts were generated by DNA synthesis (IDT gBlocks or eBlocks) or PCR (Q5 polymerase, NEB M0491S). The complete plasmid sequence for α4 and the full coding region sequences for all other plasmids are provided in .fasta format on Zenodo (https://doi.org/10.5281/zenodo.13345292). For all plasmids generated, DNA from single clones was purified (E.Z.N.A. Plasmid DNA Mini Kit I, Omega Bio-Tek D6942) and sequenced (Whole Plasmid Sequencing, Plasmidsaurus) to verify the integrity of the constructs.

### 293T-ΔNC cell culture and transfection

The 293T N-cadherin knock-out cell line (293T-ΔNC) (52) was provided by Joshua Sanes’s lab. Cells were cultured in the adherent state in a 1:1 mixture of Dulbecco’s Modified Eagle’s Medium and Ham’s F-12 medium containing L-glutamine and without phenol red (DMEM/F-12; Corning, 16-405-CV) supplemented with 10% fetal bovine serum (FBS; Corning, 35-011-CV) and 1X non-essential amino acids (NEAA; Lonza, 13-114E). Cells were maintained in exponential growth by passaging every 3-4 days. A day before transfection, 293T-ΔNC cells were seeded at 100,000 cells/well in a total volume of 400 µL medium in glass-bottom culture chambers (Cellvis, C8-1.5-N). The cells adhered to the culture chambers overnight in an incubator maintained at 37°C with 5% CO_2_.

Stocks of DNA for 293T-ΔNC transfections were prepared by diluting plasmids to 50 ng/µL with DNA elution buffer (10 mM Tris-HCl pH 8.5; Omega Bio-Tek, D6942-02) so that each transfection reaction mix contained the same amount of DNA and salts. Transfection was carried out using LipoFectMax 3000 Transfection Reagent (ABP Biosciences, FP318). For each sample, one well of cells was transfected with 0.3 µg DNA in 25 µL total volume of transfection mixture containing a 1:3:2 mass ratio of DNA, Component A, and Component B in Opti-MEM I Reduced Serum Medium (Opti-MEM; Gibco, 31985-70) per transfection reaction. The transfection reaction mixtures were incubated at room temperature for 10-15 minutes, then added directly to the medium of the cells in the culture chambers. Transfection proceeded overnight in a 37°C, 5% CO_2_ incubator. The seeded 293T-ΔNC cells remained attached to the glass substrate throughout the transfection process.

### Sample preparation and staining

Live cells were stained 20-24 hours post-transfection, and the 293T-ΔNC cells remained adhered to the glass substrate throughout staining and imaging. A FITC-conjugated antibody recognizing the Myc epitope tag (Miltenyi Biotec, 130-116-485) was diluted 1:50 in fresh medium. Three hundred and fifty microliters of culture medium were removed from each well and replaced with 50 µL diluted antibody solution (final antibody working concentration: 1:100 dilution). Unstained samples were treated identically, except that the spent culture medium was replaced with 50 µL fresh medium instead of antibody solution. Samples were incubated at 37°C, 5% CO_2_ for one hour, then 300 µL fresh medium were added to each well to dilute the antibody. Two sets of three washes were subsequently performed, with a 30-minute incubation at 37°C, 5% CO_2_ between the two sets of washes. For each wash, 200 µL medium were removed from each well and replaced with 200 µL fresh medium (antibody diluted 2X per wash; final antibody dilution 256X after all washes). Cells were imaged live immediately following the final wash.

### Microscopy

Samples were imaged using a Zeiss LSM 880 laser scanning confocal microscope using a Plan-Apochromat 10x/0.45 NA air objective. The sample was brought into focus and the pinhole was set to correspond to a 16 µm optical section, or approximately the height of a cell. Three fields of view were manually defined using only the brightfield channel for each sample well, and the fields of view were acquired sequentially using automated stage positioning with hardware autofocus to maintain the same focal plane at each position. Images were acquired in bidirectional scanning mode with a 1 µsec pixel dwell time and 4x line-by-line averaging. The two fluorescence channels were acquired sequentially on a PMT detector, and the brightfield channel was acquired in transmission mode. For the transfection marker mCherry (FP) channel, we used a 561 nm excitation laser and collected emitted light of wavelengths 578-696 nm. For the FITC-conjugated Myc antibody (Ab) channel, we used a 488 nm excitation laser and collected emitted light of wavelengths 493-592 nm.

### Image processing and quantification

Image processing was performed in Python 3 using czifile v.2019.7.2 to load the images and metadata. Cells were segmented in the brightfield channel with cellpose v.2.2.3 (69) using a GPU on the FASRC Cannon cluster supported by the FAS Division of Science Research Computing Group at Harvard University. For both fluorescence channels, the background was estimated for each individual image as the median intensity of all non-cell pixels and subtracted. *FP_cell_* and *Ab_cell_* were calculated with the single_cell.average function in microutil v.0.4.0 (70). Cells containing any saturated pixels were excluded from downstream analysis. The transfection threshold was set as the 0.995^th^ quantile of the pooled *FP_cell_* values from the mock-transfected samples in all biological replicate experiments. Thus, the probability that a cell that we designated as transfected (having an *FP_cell_* value greater than the transfection threshold) was actually not transfected was less than 0.005. The surface stain threshold was calculated similarly, as the 0.995^th^ quantile of pooled *Ab_cell_* values from samples transfected with untagged γC3 or untagged γA11. Histograms were calculated and displayed after applying a logicle transform (71) using FlowKit v.1.0.1 (72) and seaborn v.0.13.0 (73). For each sample, *f* ^+^ was calculated as the fraction of transfected cells with *Ab_cell_* values higher than the surface stain threshold, and *Ab^-^* was calculated as the average of all transfected cell *Ab_cell_* values. To calculate *E_surface_*, log-spaced bins were defined using the minimum and maximum *FP_cell_* values for all transfected cells (across biological replicate experiments and samples). Cells were assigned to bins based on their *FP_cell_* values, and *FP_bin_* and *Ab_bin_* were then calculated individually for each sample as the mean *FP_cell_* and *Ab_cell_* values per bin, respectively. *E_surface_* was calculated as the slope of the least-squares linear regression of the log of *FP_bin_* and *Ab_bin_* using the linregress function in scipy (68), excluding bins with negative *Ab_bin_* values (this only occurred for a few bins in untagged isoform samples). Of note, the data provide single-cell measurements for *FP_cell_* and *Ab_cell_*, so the linear regression and slope can be computed either with or without binning. We chose to use bins as they reduce the contribution of the many transfected cells with very low expression levels. P-values were calculated using the statistical tests indicated in the text or figure legends.

### K562 cell culture, electroporation, and staining

K562 cells were provided by Daniel A. Fletcher’s lab and the UC Berkeley Cell Culture Facility and were maintained in suspension between 1×10^5^ and 1×10^6^ cells/mL in Roswell Park Memorial Institute 1640 medium (RPMI; Corning 10-040-CV) supplemented with 10% FBS and 1 mM sodium pyruvate (Gibco 11360070). Twenty µL of cells at 1×10^7^ cells/mL were electroporated with two µL of DNA at 500–1000 ng/µL concentration with two 10-ms width, 1450 V pulses using a Neon electroporation system and a 10 µL Neon tip (Invitrogen MPK1025). After electroporation, cells recovered for 18-24 hours in complete medium at 37°C with 5% CO_2_. For staining, cells were centrifuged at 500 × g for 5 minutes and resuspended in complete medium containing the anti-Myc antibody diluted 1:100. After incubating for one hour at 37°C, 5% CO_2_, the cells were centrifuged and resuspended in complete medium without antibody three times to wash. Cells were analyzed on an Attune CytPix Flow Cytometer, and surface delivery quantification was performed as described for imaging data.

## Acknowledgements

The authors thank Barry Honig, Tom Maniatis, and Erin Flaherty for supplying plasmids, Josh Sanes for supplying 293T-ΔNC cells, Daniel A. Fletcher for supplying K562 cells, and John Russell, José Velilla, and Sam Berry for helpful feedback on the manuscript. We also thank Brenda Chiang and Nhu Dang for early work on the project and the Harvard Center for Biological Imaging (RRID:SCR_018673) for infrastructure and support. R.G. acknowledges funding from a Harvard Brain Science Initiative Bipolar Disorder Seed Grant and NIH grant R01GM120996. E.J.M. thanks the NSF-Simons Center for Mathematical and Statistical Analysis of Biology at Harvard, award number #1764269 and the Harvard Quantitative Biology Initiative, Harvard Physics of Living Systems, the Aramont Fund for Emerging Science Research, and the Simmons family for generous funding.

## Data availability

Microscopy datasets, plasmid sequences, alignment files, and raw data files containing values for the figures in the manuscript are available on Zenodo (https://doi.org/10.5281/zenodo.13345292).

Python scripts are available on GitHub (https://github.com/emay2022/surface-trafficking).

Plasmids are available from the authors upon reasonable request.

