## Supplemental Figures for "Surface delivery quantification reveals distinct trafficking efficiencies among clustered protocadherin isoforms"

**This PDF file includes:**

Figures S1-S6

Table S1

**Other supporting materials for this manuscript include the following:**

Dataset S1

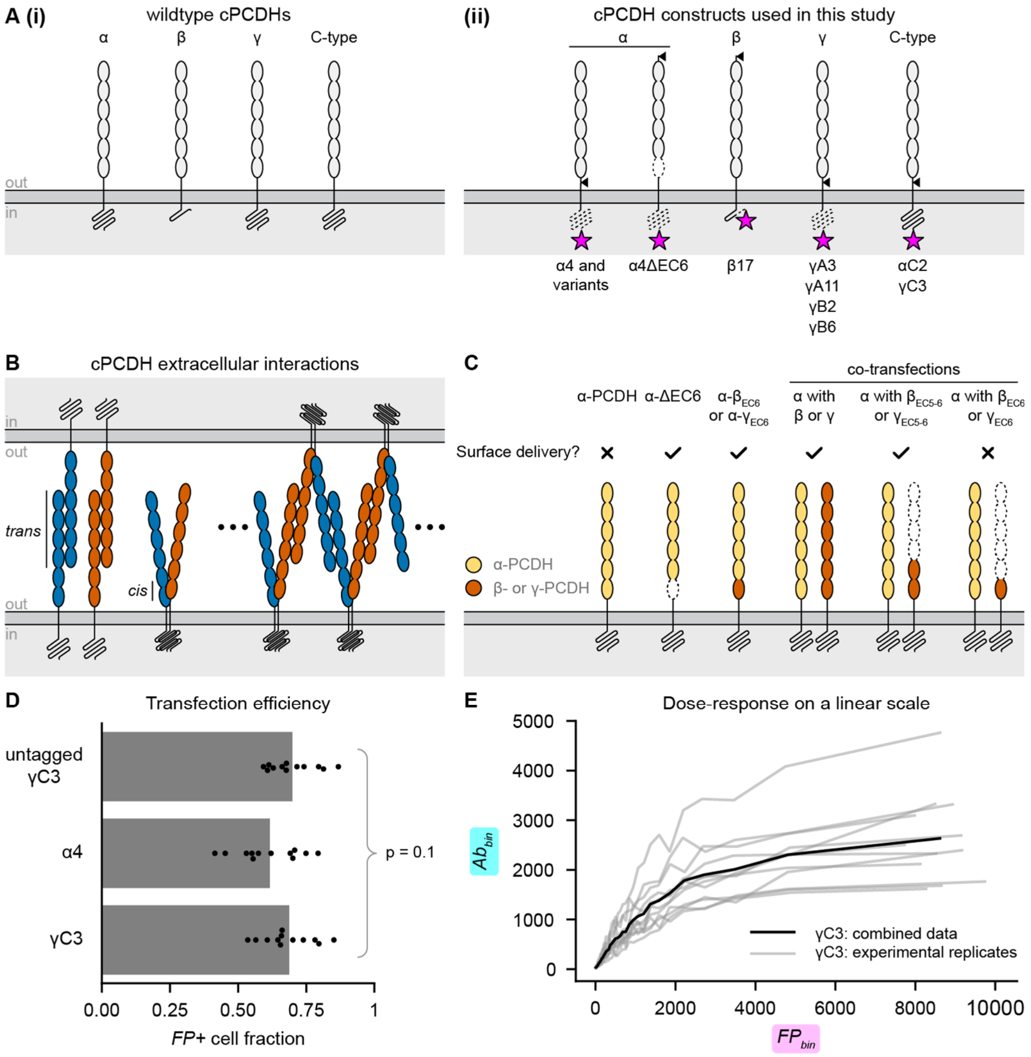


**Figure S1.** Schematics of cPCDHs and their functions in neuronal self-avoidance and surface trafficking. (A) Part (i) shows schematics of wildtype cPCDH proteins. Part (ii) shows schematics of the cPCDH constructs used in this study. The black triangle indicates the position of the Myc tag, the magenta star indicates the C-terminal fluorescent protein, and dotted lines indicate deleted portions of the sequence as compared to wildtype. (B) Diagrams of cPCDH extracellular interactions, with isoforms represented as different colors. The *trans* interaction involves EC1-4 and is strictly homophilic. The *cis* interaction is preferentially heterophilic and asymmetric, involving EC5-6 of one subunit and only EC6 of the partner subunit. The combination of *cis* and *trans* interactions can lead to extended cPCDH assemblies. (C) Schematic summary of previous experiments from Thu and colleagues using cell aggregation assays and surface staining with microscopy in K562 cells to identify perturbations that lead to α-PCDH surface delivery, with the reported binary yes-or-no readout indicated as check and cross symbols, respectively (1). (D) Transfection efficiency for untagged γC3, α4, and γC3. The p-value was calculated using a one-way ANOVA. (E) $Ab_{bin}$ versus $FP_{bin}$ for $E_{surface}$ calculations for the γC3 sample in Figure 2, shown on a linear scale. Individual experimental replicates (gray lines) and combined data over all experimental replicates (black line) are shown.

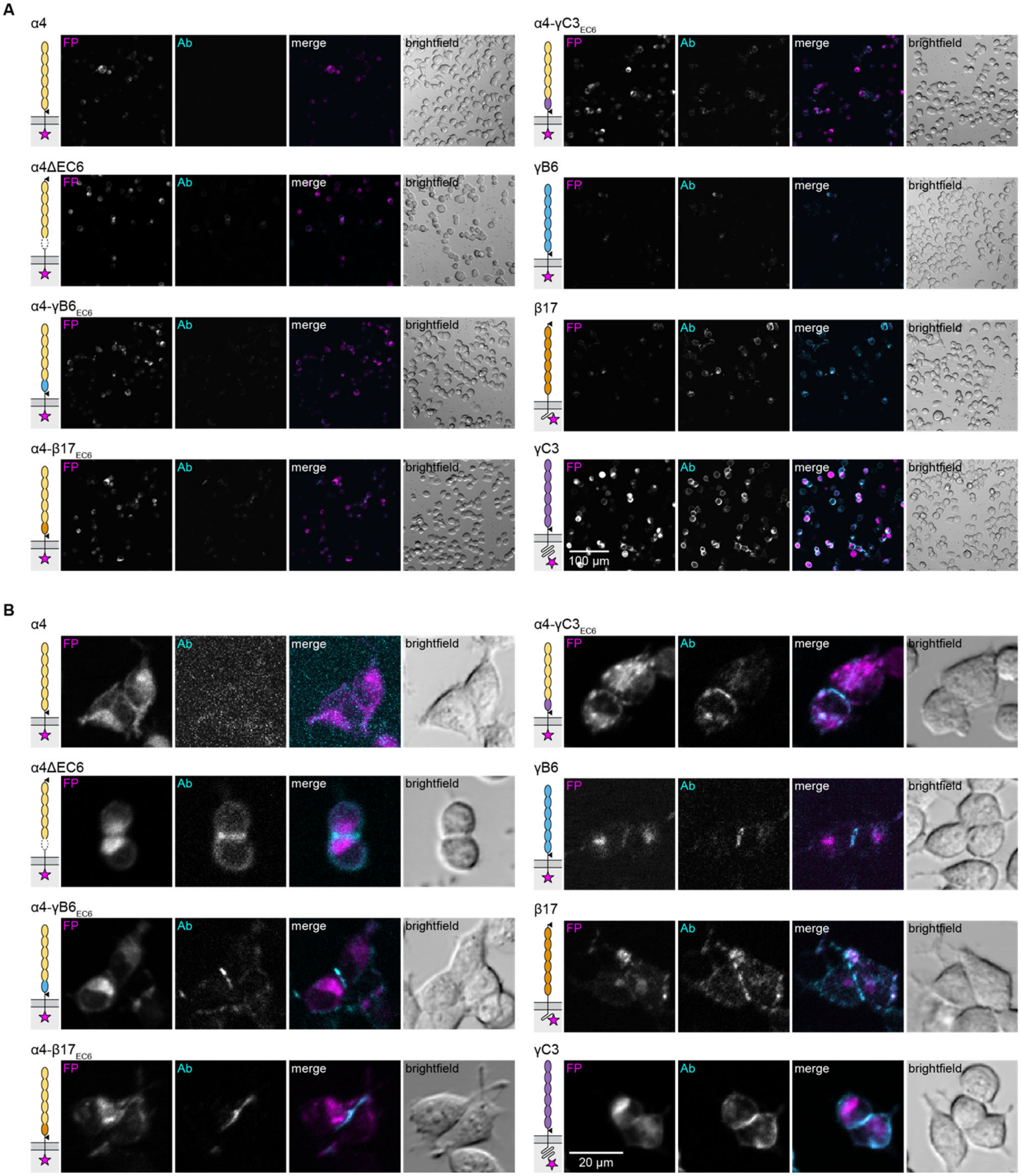


**Figure S2.** Example images from surface staining experiments. Individual fluorescence channels are shown in grayscale and a merged image is shown with the FP channel in magenta and the Ab channel in cyan. (A) Overview images including many cells per field of view. All fluorescence images were contrast-adjusted to the same limits. Scale bar, 100 µm. (B) Cropped regions featuring transfected cell pairs highlight the presence of extracellular cPCDH staining at cell-cell junctions for all surface-trafficking constructs, but not for α4. Contrast limits were adjusted individually for each image to best display the junctional staining. Scale bar, 20 µm.

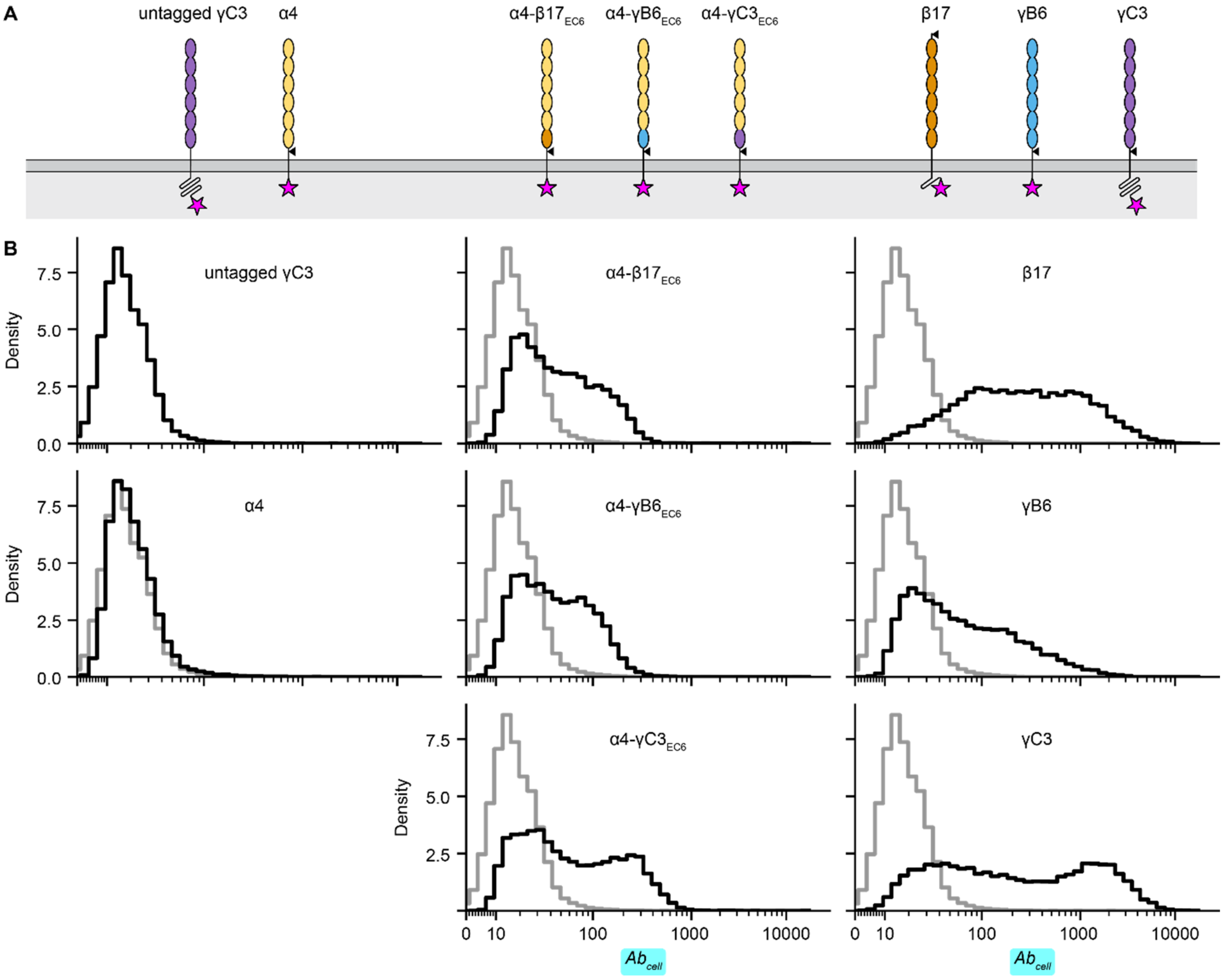


**Figure S3.** Even surface-localizing isoform samples contain a population of transfected cells that remain unstained. (A) Construct schematics. (B) $Ab_{cell}$ histograms for each of the constructs shown in A. The histogram for the untagged γC3 construct (negative control for Ab staining) is reproduced in grey on all graphs as a reference.

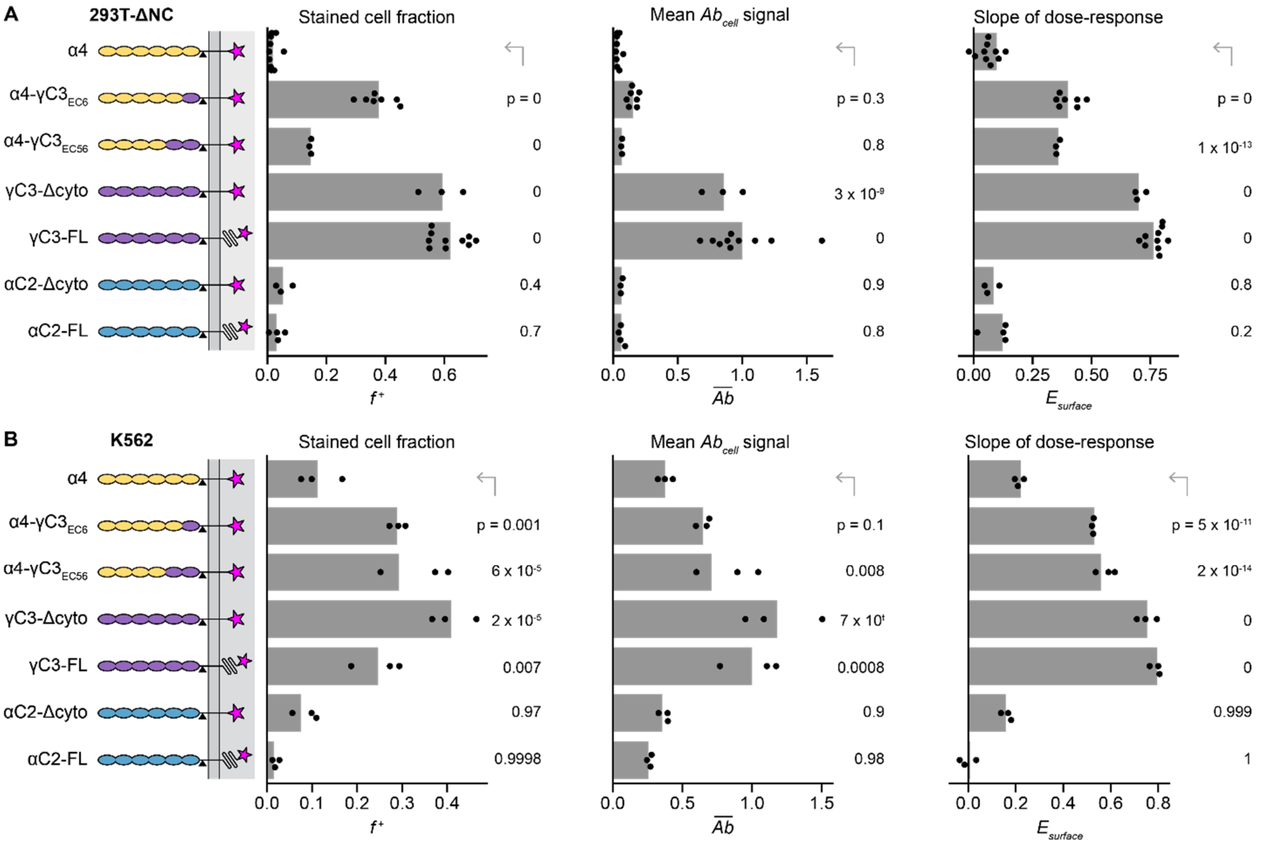


**Figure S4**. Surface trafficking measurements in 293T-ΔNC cells using microscopy match those observed in K562 cells using flow cytometry. (A) Quantification of $f^+$, $\overline{Ab}$, and $E_{surface}$ for the constructs shown using 293T-ΔNC cells imaged with fluorescence microcopy. (B) Quantification of $f^+$, $\overline{Ab}$, and $E_{surface}$ for the same constructs using K562 cells measured with flow cytometry. $\overline{Ab}$ values are normalized to the mean $\overline{Ab}$ of the positive control construct, full-length Myc-tagged γC3 (γC3-FL). Points are biological replicates ($n$ = 3 for α4-γC3EC5-6, γC3Δcyto, and αC2Δctyo; $n$ = 4 for αC2-FL; $n$ = 7 for α4-γC3EC6; $n$ = 10 for α4 and γC3-FL in panel A); $n$ = 3 for all samples in panel B), and bars are mean values across replicates. The p-values for each sample compared to α4 were calculated using a one-tailed Dunnett's test.

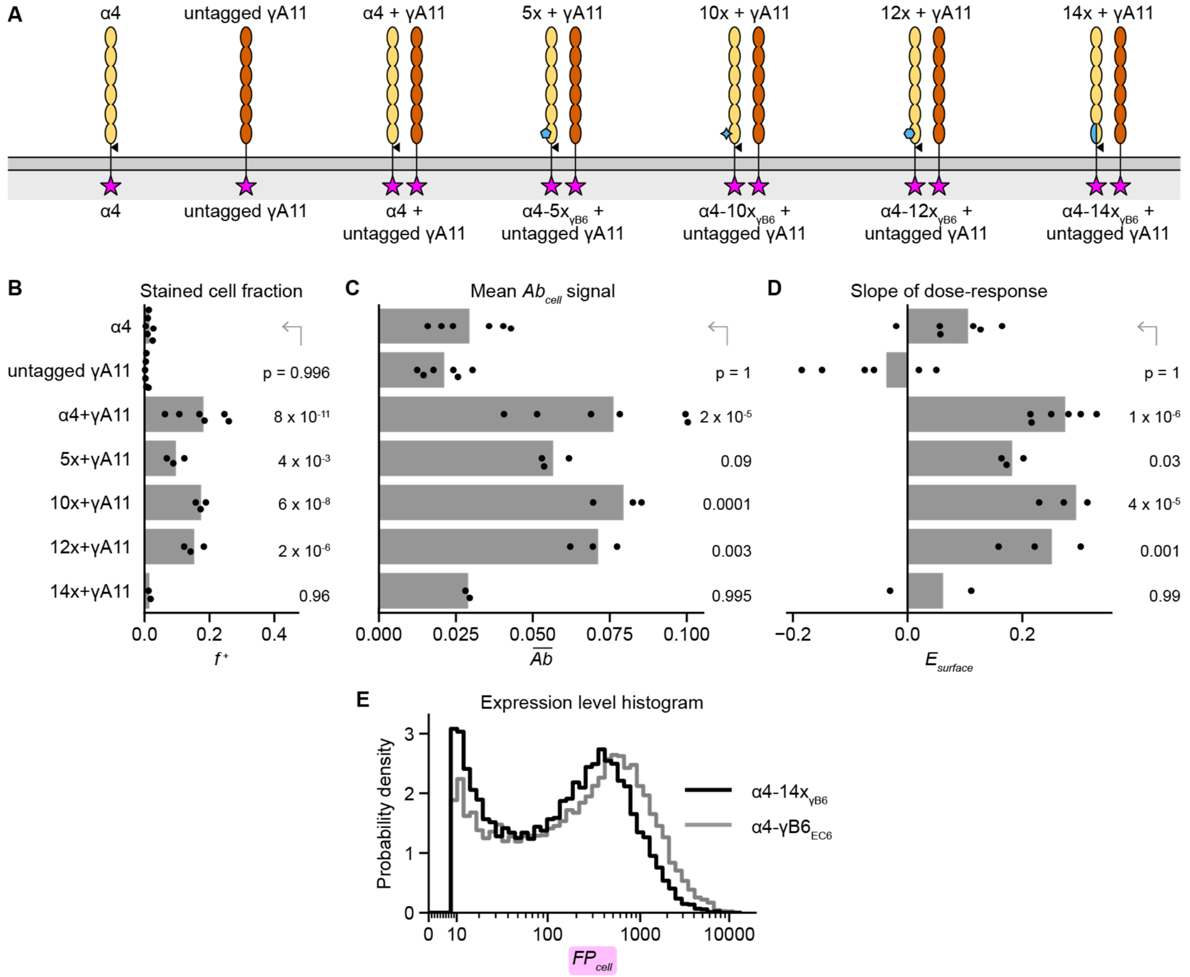


**Figure S5.** Co-transfection experiments report on the stability of chimeric α4 variants. (A) Schematics of constructs transfected together in co-transfection experiments. In each case, only the α4 variant has a Myc tag, so Ab signal only reports on the presence of the α4 variant on the cell surface. Untagged γA11 is included as a control for Ab specificity, α4 transfected alone is included as the negative control, and α4+γA11 is included as a positive control. (B-D) Quantification of $f^{+}$ (B), $\overline{Ab}$ (C), and $E_{surface}$ (D) for the experiments schematized in panel A. In C, $\overline{Ab}$ values are normalized to the mean $\overline{Ab}$ of the positive control α4-γB6$_{EC6}$. Points are biological replicates ($n$ = 6 for α4, untagged γA11, and α4+γA11; 3 for 5x+γA11, 10x+γA11, and 12x+γA11; and 2 for 14x+γA11), and bars are replicate means. The p-values for each sample compared to α4 were calculated using a one-tailed Dunnett's test. (E) Expression level histograms show the distribution of $FP_{cell}$ values for α4-14x$_{γB6}$ (black line) overlayed with α4-γB6$_{EC6}$ (gray line).

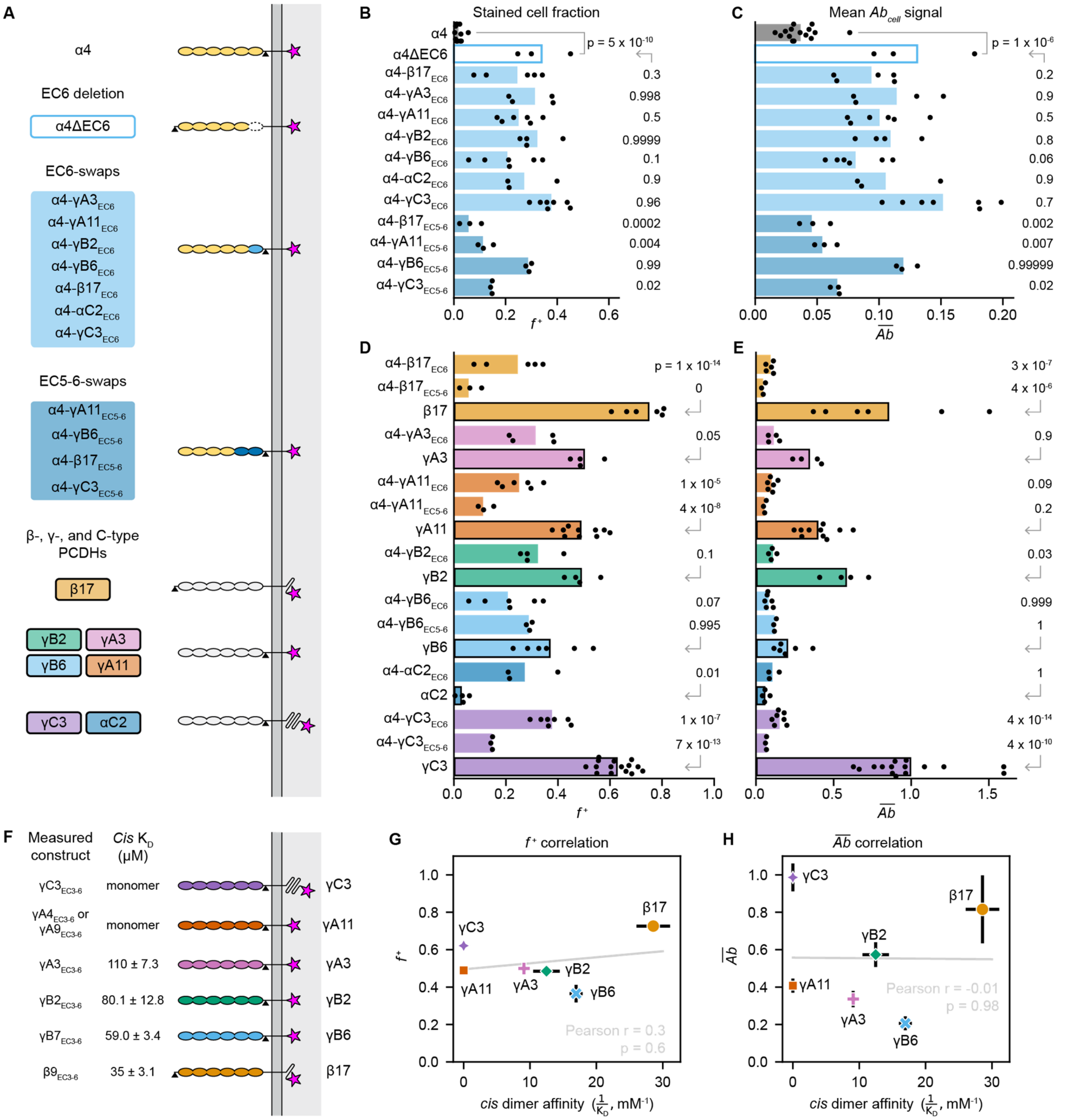


**Figure S6.** Graphs of $f^+$ and $\overline{Ab}$ supporting Figure 4. (A) Detailed construct schematic and color coding for variants shown in panels B-G and in Figure 4. (B-C) Graphs comparing $f^+$ (B) and $\overline{Ab}$ (C) for α4 variants with EC6 deleted (unfilled lighter blue bar), EC6 swapped (lighter blue bars), and EC5-6 swapped (darker blue bars) to wildtype α4 (gray bars). Points are biological replicates ($n$ = 15 for α4 and γC3; 3 for α4ΔEC6, α4-αC2EC6, α4-β17EC5-6, α4-γA11EC5-6, α4-γB6EC5-6, and γC3EC5-6; 4 for α4-γA3EC6 and α4-γB2EC6; 5 for α4-β17EC6; 6 for α4-γA11EC6 and α4-γB6EC6; and 7 for α4-γC3EC6), and bars are replicate means. The p-value for α4ΔEC6 compared to α4 was calculated using a one-tailed Welch's t-test, and p-values for other samples compared to α4ΔEC6 were calculated using Dunnett's test. (D-E) Graphs of $f^+$ (D) and $\overline{Ab}$ (E) comparing constructs paired by isoform. Bars are color-coded by isoform. Bars without borders are α4 variants (EC6 or EC5-6 swaps). Bars with black borders are wildtype cPCDHs. Points are biological replicates (*n* = 15 for α4 and γC3; 3 for α4-αC2$_{EC6}$, α4-β17$_{EC5-6}$, α4-γA11$_{EC5-6}$, γB6, α4-γB6$_{EC5-6}$, and γC3$_{EC5-6}$; 4 for αC2, γA3, α4-γA3$_{EC6}$, γB2, and α4-γB2$_{EC6}$; 5 for α4-β17$_{EC6}$; 6 for β17, α4-γA11$_{EC6}$, γB6, and α4-γB6$_{EC6}$; 7 for α4-γC3$_{EC6}$; and 10 for γA11), and bars are replicate means. The p-values for each α4 variant compared to the corresponding wildtype isoform were calculated using Tukey's test. (F) Schematics of wildtype cPCDHs and reported solution affinity measurements of purified *cis*-interacting fragments (or their closest paralog) (2-5). Binding affinity is reported as $K_D$, which is high for weak interactions and low for strong interactions. (G-H) Scatter plots of β- and γ-PCDH *cis* dimerization affinity versus the surface delivery metrics $f^+$ (G) and $\overline{Ab}$ (H). Affinity is plotted as $1/K_D$ (i.e., $K_A$), with monomeric isoforms given an affinity of zero. The Pearson r statistic and its associated p-value indicate the degree of correlation.

**Table S1. Isoforms and variants used and the cPCDH subfamilies they represent**

| Subfamily | Isoform | EC6-swapped α4 variant | EC56-swapped α4 variant |
|---|---|---|---|
| α | α4* | α4ΔEC6[†] | - |
| β | β17[†] | α4-β17$_{EC6}$[†] | α4-β17$_{EC5\text{-}6}$ |
| γA | γA3 | α4-γA3$_{EC6}$ | - |
| | γA11 | α4-γA11$_{EC6}$ | α4-γA11$_{EC5\text{-}6}$ |
| γB | γB2 | α4-γB2$_{EC6}$ | - |
| | γB6[†] | α4-γB6$_{EC6}$[†] | α4-γB6$_{EC5\text{-}6}$ |
| C-type isoforms | αC2 | α4-αC2$_{EC6}$ | - |
| (polyphyletic group) | γC3* | α4-γC3$_{EC6}$[†] | α4-γC3$_{EC5\text{-}6}$ |

| Chimeric α4 mutations | Residues mutated from α4 to γB6 |
|---|---|
| α4-5x$_{\gamma B6}$ | E567Y, H589Y, R595V, Y603H, Y627R |
| α4-10x$_{\gamma B6}$ | 5x plus T564R, V577L, S578F, V590L, S632R |
| a4-12x$_{\gamma B6}$ | 10x plus E569A, A592T |
| a4-14x$_{\gamma B6}$ | 12x plus R579D, L580M |

* When included in subsequent figures, data reproduced from **Figure 2**

[†] When included in subsequent figures, data reproduced from **Figure 3**

**Dataset S1 (separate file).** Tukey's test p-values calculated for all figures in the manuscript that compare more than three samples.
